# Abnormalities of placental development and function are associated with the different fetal growth patterns of hypoplastic left heart syndrome and transposition of the great arteries

**DOI:** 10.1101/388074

**Authors:** Weston Troja, Kathryn J. Owens, Jennifer Courtney, Andrea C. Hinton, Robert B. Hinton, James F. Cnota, Helen N. Jones

## Abstract

**Background:** Birthweight is a critical predictor of congenital heart disease (CHD) surgical outcomes. Hypoplastic left heart syndrome (HLHS) is cyanotic CHD with known fetal growth restriction and placental abnormalities. Transposition of the great arteries (TGA) is cyanotic CHD with normal fetal growth. Comparison of the placenta in these diagnoses may provide insights on the fetal growth abnormality of CHD.

**Methods:** Clinical data and placental histology from placentas associated with Transposition of the Great Arteries (TGA) were analyzed for gross pathology, morphology, maturity and vascularity and compared to both control and previously analyzed HLHS placentas [1]. RNA was isolated from HLHS, TGA and control placentas and sequenced by Illumina HiSeq.Gene, analysis was performed using TopHat, R and MSigDB. Cluster analysis was performed using GoElite and Pathway analysis performed using PANTHERdb Overrepresentation Test. Immunohistochemistry was utilized to assess placental nutrient transporter expression in all three groups.

**Results:** Placental weight was reduced in TGA cases, and demonstrated reduced villous vasculature, immature terminal villi, and increased fibrin deposition in the parenchyma compared to controls and reflected our previous data from HLHS placentas. However, birth weight was not reduced in TGA cases compared to controls in contrast to the HLHS cohort and birthweight:placental weight ratio was significantly increased in TGA cases but not HLHS compared to control. Need to include RNA and IHC.

**Conclusions:** Despite common vascular disturbances in placentas from HLHAs and TGA, these do not account for the

## Introduction

Congenital heart disease (CHD) affects nearly 1% of annual births and is the leading cause of infant mortality related to birth defects. CHD is further complicated by high incidence of fetal growth restriction and prematurity. These perinatal outcomes strongly impact neonatal cardiac surgical mortality and morbidity but also adversely impact long-term functional status, quality of life and healthcare costs [2-4]. The etiology of fetal growth abnormalities in CHD is largely unknown.

The placenta plays a central role in fetal development, regulating the transport of nutrients and oxygen from mother to the fetus [5] and, therefore, is a good target for investigating growth disturbance in fetal CHD. Placental abnormalities are common in prenatal disorders and estimated to cause the majority of fetal growth restriction. Recent studies have demonstrated that placental complications and impaired fetal growth are associated with CHD [6-9]. The rate of fetal nutrient uptake and oxygen diffusion is influenced by placental size, vascular surface area, transporter capacity, membrane permeability and thickness, and rate of uterine blood flow. Placental villi grow and remodel throughout gestation, increasing the villous surface area needed for optimal transport. By term, the villi mature into an expansive network of terminal villi to support the accelerated growth of the fetus. In addition to these morphologic changes, both placenta and fetus require adequate transport of amino acids, fatty acids, and glucose from the maternal circulation to achieve proper growth[5]. The delivery of total amino acids from mother to fetus is dependent upon the placental transport capacity and strongly correlated with the rate of fetal growth[10]. Placental expression and activity of amino acid transport system A (isoform SNAT) and L (LAT1/2) are markedly reduced in animal models and cases of human fetal growth restriction [11]. Similarly, alterations to the expression of placental glucose transporter isoforms, GLUT1 and GLUT3, have been implicated in pathological pregnancies and diminished fetal growth[12] Disruption of this process can negatively impact fetal growth and development but has not been explored in the setting of fetal CHD.

We previously identified altered placental vasculature, placental insufficiency, and morphologic abnormalities in placentas from cases of hypoplastic left heart syndrome (HLHS), a form of CHD associated with significant growth disturbances [1]. To build on those findings, we performed a similar analysis in cases of transposition of the great arteries (TGA), a form of CHD with normal birth weights, comparing the findings. Furthermore, the morphologic assessment was expanded to include transcriptome and transporter analysis HLHS, TGA, and controls. Given the known differences in birth weight, we hypothesized TGA placentas would have a greater number of terminal villi, more vessels, and higher transporter function when compared to HLHS placentas..

## Methods

The cohort of patients was assembled retrospectively from a single center case series at Cincinnati Children’s Hospital Medical Center (CCHMC) and Good Samaritan Hospital (GSH, Cincinnati, OH) in 2003-2014. Maternal, fetal, and neonatal clinical data were collected at full term (>37 weeks gestation) from neonates with either HLHS or TGA. Cases with genetic complications, multiple gestation pregnancies, history of maternal diabetes, preeclampsia, or hypertension were excluded. TGA was defined as having the aorta arise predominantly from the right ventricle and pulmonary artery from the left ventricle. HLHS were defined as previously published [1]. Controls were defined as full term births with no maternal or fetal abnormalities. The Institutional Review Boards of CCHMC and GSH approved this study.

### Clinical data

Medical record review was performed for the collection of clinical pregnancy data, birth anthropometrics and gross pathology assessment.

### Placental morphology

Placental tissues were cut into 5µm sections and serial sections were deparaffinized in xylene and rehydrated. Sections were stained with Hematoxylin and Eosin (H&E) and assessed for evidence of inflammation and alterations in villous architecture. To assess the terminal branching of placental villi that normally occurs exponentially in the third trimester, terminal villi (40–80µm in diameter in normal placentas) density in each high-powered field was assessed both qualitatively and the number of villi less than 80µm in diameter counted. To investigate the villous vasculature, the number of vessels per high-powered field was assessed initially by manual counting in 5 random high-powered fields across 3 sections following immunohistochemistry for CD-31. Vessel density is represented as the average count of vessels per high-powered field and the range of counts. To aid in assessment of maturation, vasculo-syncytial counts were performed as previously described [1]

### Isolation and analysis of RNA from FFPE samples

RNA was extracted from formalin-fixed, paraffin-embedded (FFPE) human placenta tissue blocks using the RecoverAll Total Nucleic Acid Isolation kit (Ambion). Three-20um sections were cut from the center of the FFPE blocks and RNA was extracted following manufacturer’s instructions using a 2 hour proteinase K digestion at 50°C. RNA integrity was assessed by DV200 using a Fragment Analyzer (Advanced Analytical), specifically to determine the applicability of degraded RNAs to NGS library preparation.

### Target RNA enrichment

Ribo-Zero Gold kit (Illumina, San Diego, CA) was used to deplete rRNA before library preparation for RNA-seq. Ribo-Depletion script was run on WaferGen (Fremont, CA) Apollo 324 system to automate RNA depletion with 1 µg of total RNA as input.

### RNA-seq library preparation

The RNAseq library was prepared by using NEBNext Ultra Directional RNA Library Prep kit (New England BioLabs, Ipswich, MA). During the second cDNA synthesis dUTP was incorporated into the second cDNA strand to maintain strand specificity. The purified cDNA was end repaired, dA tailed, and ligated to an adapter with a stem-loop structure. The dUTP-labelled 2nd strand cDNA was removed by USER enzyme to maintain strand specificity. The libraries were amplified with PCR and purified alongside the library prep negative controls via AMPure XP beads for QC analysis. The quality and yield of the library were analyzed by Bioanalyzer (Agilent, Santa Clara, CA) using DNA high sensitivity chip. To accurately quantify the library concentration for the clustering, the library was diluted 1:104 in dilution buffer (10 mM Tris-HCl, pH 8.0 with 0.05% Tween 20) and measured by NEBNext Library Quant Kit (New England BioLabs) using QuantStudio 5 Real-Time PCR Systems (Thermo Fisher, Waltham, MA).

### HiSeq Sequencing

To study differential gene expression, individually indexed and compatible libraries were proportionally pooled (∼50 million reads per sample in general) for clustering in cBot system (Illumina, San Diego, CA). Libraries at the final concentration of 15 pM were clustered onto a single read (SR) flow cell using Illumina TruSeq SR Cluster kit v3, and sequenced to 50 bp using TruSeq SBS kit on Illumina HiSeq system.

### mRNA-sequence data analyses

Adapters were trimmed from the reads using CutAdapt V1.8.1[13]. TopHat V 2.0.13 [14] and Bowtie V2.1.0[15] were used to align the reads to Human Genome GRCh37 (hg19) and generate BAM files for further analyses. AltAnalyze [16] was used to determine differential gene expression between the sample groups, HLHS Vs. term, HLHS Vs. TGA and TGA Vs. term. Differentially expressed genes were clustered using GO-Elite with a calculated Z-score of 1.96 and p < 0.05 using Fisher Exact Test [17]. Brifely, unbiased clustering was performed by AltAnalyze (V 2.1.0) using the clustering algorithm HOPACH. Following alignment of read counts using TopHat and pruning with an RPKM cutoff of 1 million reads, pairwise group comparisons were generated between all possible HOPACH clusters. Further pruning to include only protein-encoding genes was performed, followed by heat map and gene list generation.

### Pathway analysis

Utilizing Panther db [18, 19], we performed pathway analysis and investigated the over-representation of biological processes in the lists of upregulated or downregulated genes seen between HLHS and control or TGA placentas (Use the Bonferroni correction for multiple testing PANTHER Overrepresentation Test (release 20170413). A q value of 0.25 was utilized to generate gene lists for pathway analysis). Gene Ontology and Reactome databases were utilized for analysis of gene lists relative to molecular functions, biological processes or pathways.

### Immunohistochemistry

For antibody staining, sections were incubated in 1X sodium citrate (pH 6.0) target retrieval solution (Dako Cytomation, Carpathia, CA) at 95°C for 30 minutes, followed by a 20 minute cool down at room temperature. After washing, sections were incubated in 3% hydrogen peroxide for 10 minutes and subsequently washed in PBS with 1% tween. Nonspecific binding was blocked by incubation with 5% normal goat serum and 1% BSA for 1 hour at room temperature. Sections were incubated with primary antibody in diluent overnight at 4°C. With the exception of CD98 (Abcam, ab23495, 1:100), which was grown in mouse, all primary antibodies were derived from rabbit serum: anti-CD31 (Abcam, ab28364), anti-Leptin (LSBio, LS-B1459, 1:250), anti-GLUT1 (Abcam, ab15309, 1:750), anti-GLUT3 (Abcam, ab15311, 1:150), anti-SNAT2 (LSBio, LS-C179270, 1:500), anti-LAT1 (Abcam, ab8526, 1:500), anti-LAT2 (Abcam, ab75610, 1:250), and anti-TAUT (LSBio, LS-C179237). After incubation, sections were washed and incubated with appropriate secondary antibodies in diluent (Vector Laboratories, Inc., Burlingame, CA, 1:1000) at room temperature for 1 hour except anti-SNAT2 was incubated for 2 hours. Target antigen signal was amplified with ABC (Vector Laboratories) and antibody binding was detected using DAB (Dako Cytomation). Slides then were counterstained with hematoxylin, dehydrated, and mounted. Histologic examination and analysis were conducted by light microscopy using a Nikon Eclipse 80i microscope.

### Statistical Analysis

Data were explored using tests of normality and evaluation of skew. Normally distributed data are expressed as mean ± standard deviation (SD) and analyzed with Student’s T-test and ANOVA as appropriate. Tukey-Kramer Multiple Comparisons Test were included as Posthoc tests if ANOVA reached significance. Non-normally distributed data are expressed as median and range and analyzed by Mann Whitney U test.

## Results

### Maternal & Fetal demographics

There were no significant differences in maternal age, BMI or gestational age between TGA, HLHS and control cases. Birthweight and placental weight of the TGA group was similar to the control group. However, birth head circumference of TGA patients were significantly lower than controls(p < 0.001, Table 1), consistent with previous findings [20, 21]. Interestingly, there was a higher proportion of male to female offspring (6:2) in the TGA cohort compared to control (7:11). There were no cases of SGA in either the TGA or control cohort. The ratio of birthweight to placental weight was significantly (ANOVA, p=0.004, n>8 per group) increased in the TGA (7.04± 0.34), but not HLHS (6.69±0.46) group, compared to control (6.21± 0.63). Interestingly, posthoc testing demonstrated no significant difference in birthweight: placenta weight ratio between the TGA and HLHS placentas.

**Table 1:**
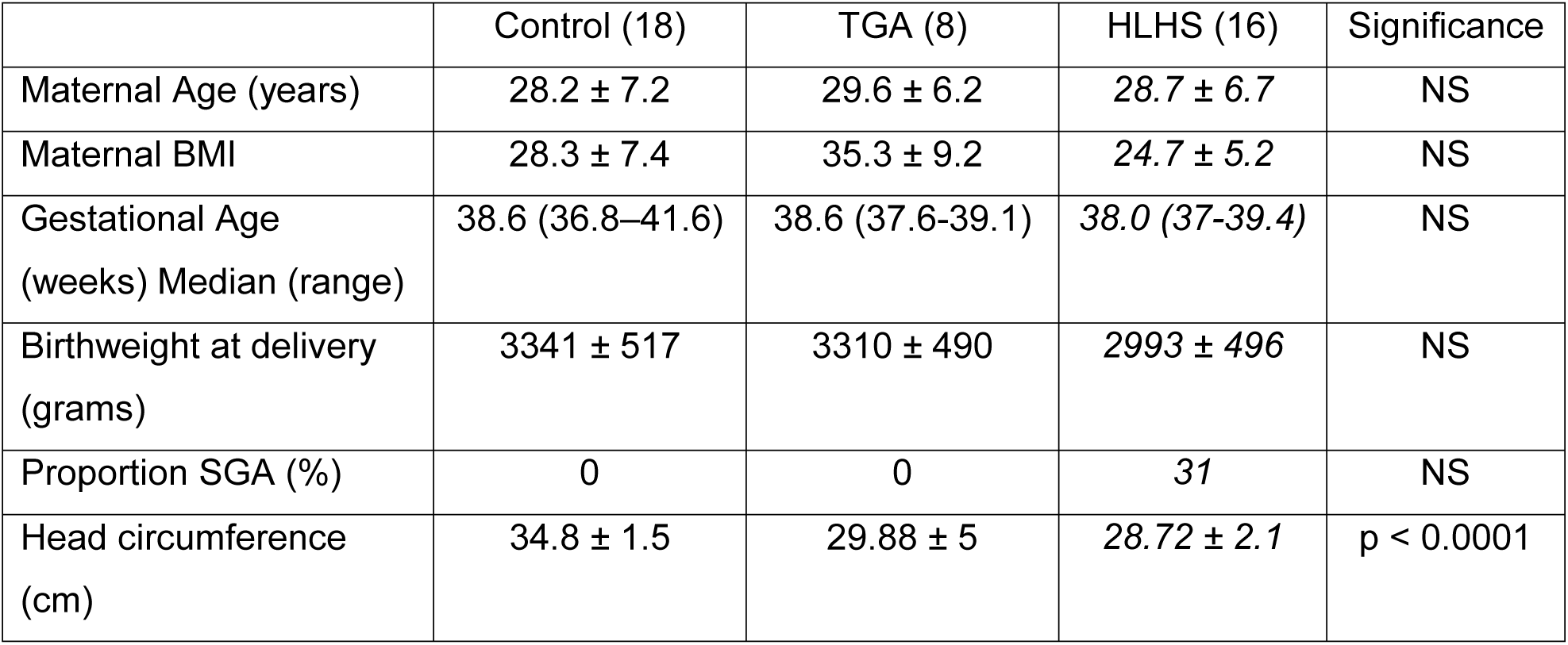
Maternal and fetal characteristics. Data are presented as mean ± standard deviation or median and range. TGA: Transposition of the Great Arteries; HLHS: Hypoplastic Left Heart Syndrome; BMI: Body Mass Index; SGA: Small for Gestational Age; NS: Not Significant.* Placental efficiency is defined as gram of fetus per gram of placenta. HLHS data in italics was previously published ([1]) but included for comparison with TGA.

|  | Control (18) | TGA (8) | HLHS (16) | Significance |
| --- | --- | --- | --- | --- |
| Maternal Age (years) | 28.2 ± 7.2 | 29.6 ± 6.2 | 28.7 ± 6.7 | NS |
| Maternal BMI | 28.3 ± 7.4 | 35.3 ± 9.2 | 24.7 ± 5.2 | NS |
| Gestational Age<br>(weeks) Median (range) | 38.6 (36.8–41.6) | 38.6 (37.6-39.1) | 38.0 (37-39.4) | NS |
| Birthweight at delivery<br>(grams) | 3341 ± 517 | 3310 ± 490 | 2993 ± 496 | NS |
| Proportion SGA (%) | 0 | 0 | 31 | NS |
| Head circumference<br>(cm) | 34.8 ± 1.5 | 29.88 ± 5 | 28.72 ± 2.1 | p < 0.0001 |

### Placental pathology and morphology

As we previously demonstrated [1] HLHS placentas have reduced villous vasculature and reduction in arborization (lower numbers of terminal villi), as well as other hallmarks of immaturity. Intervillous fibrin deposition and prominent syncytial nuclear aggregates were observed in the gross pathology of over 50% of the TGA placentas with no evidence of increased chorioamnionitis, inflammation, thrombi, or infarction. Histological analysis of the placentas revealed the TGA group had significantly reduced villous blood vessels per high-powered field when compared to controls (p<0.0001, Table 2). Similarly, the number of terminal villi per high power field and vasculo-syncytial membrane counts were significantly reduced in TGA placentas Vs. term (Table 2),

**Table 2:** Placental Characteristics. Data are presented as mean ± standard deviation or median and range. TGA: Transposition of the Great Arteries; HLHS: Hypoplastic Left Heart Syndrome. HLHS data in italics was previously published ([1]) but included for comparison with TGA.

|  | Control (18) | TGA (8) | HLHS(18) | Significance |
| --- | --- | --- | --- | --- |
| Placental weight (grams) | 538 ± 122 | 470 ± 78 | 447±104 | P<0.05 |
| Terminal villi/hpf | 21.3±3.9 | 16.3±2.4 | 10.0±3.4 | P<0.05 |
| Vessel density/hpf | 69 (51-81) | 44 (21-61) | 35 (11-62) | P<0.0001 |
| Vasculo-syncytial membrane count/hpf | 10.4 ± 2.5 | 2.68 ± 2.1 | 3.42 ± 2.27 | P<0.01 |

### Genome-wide transcriptomic analyses

Differentially expressed (DE) genes were identified between comparisons of control, HLHS, and TGA (Figure 3).

GO-Elite identified five clusters of 171, 11, 41, 51, and 11 genes each. In the largest cluster, genes involved in left/right patterning and symmetry, heart looping, muscle development, and mitochondrial energy synthesis are significantly downregulated in the HLHS placentas compared to control and TGA. Human leukocyte antigens (HLA) and transcriptional regulators were increased in the TGA placentas compared to control and HLHS (second and third clusters). Pregnancy-related genes and hormonal stimulators were significantly upregulated in the HLHS and TGA placentas in the fourth cluster. The fifth cluster included nutrient transporters, which were increased in the HLHS placentas.

### Panther Pathway analysis

Over-represented pathways and biological process identified utilizing PantherDB over-representation are presented in Table 3-6. Interestingly, there were no pathways or processes identified as significantly over-represented in the gene lists that were down-regulated in TGA versus control or HLHS.

**Table 3:** Over-representation of Reactome pathways in genes identified as down-regulated in HLHS vs control.

| Reactome pathways | Homo sapiens (REF) | Client Text Box Input ( Hierarchy ) |  |  |  |  |
| --- | --- | --- | --- | --- | --- | --- |
|  |  | # | expected | Fold Enrichment | +/- | P value |
| Generation of second messenger molecules | 37 | 7 | .35 | 20.00 | + | 1.51E-04 |
| Interferon Signaling | 192 | 10 | 1.82 | 5.51 | + | 3.17E-02 |
| RHO GTPase Effectors | 261 | 12 | 2.47 | 4.86 | + | 1.59E-02 |
| Cell Cycle, Mitotic | 464 | 18 | 4.39 | 4.10 | + | 1.02E-03 |
| Cell Cycle | 568 | 21 | 5.37 | 3.91 | + | 2.46E-04 |

**Table 4:**
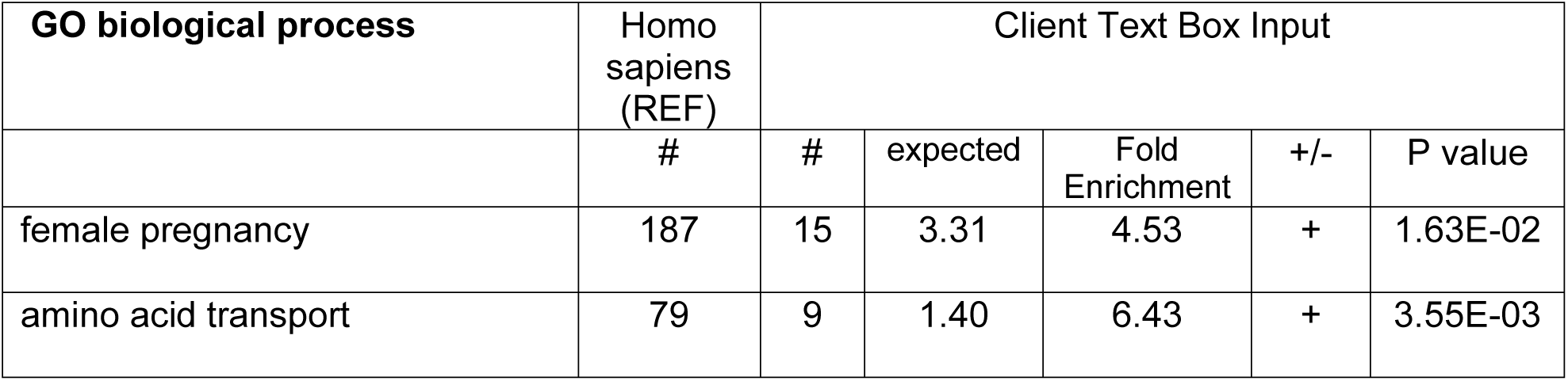
Over-representation of GO biological processes in upregulated between HLHS and control.

**Table 5:** Upregulated TGA vs HLHS. Over-representation of Reactome pathways in upregulated TGA vs HLHS

| Reactome pathways | Homo sapiens (REF) | Client Text Box Input |  |  |  |  |
| --- | --- | --- | --- | --- | --- | --- |
|  |  | # | expected | Fold Enrichment | +/- | P value |
| Respiratory electron transport. | 120 | 33 | 14.57 | 2.27 | + | 3.82E-02 |
| The citric acid (TCA) cycle | 162 | 43 | 19.66 | 2.19 | + | 5.76E-03 |
| Generic Transcription Pathway | 833 | 146 | 101.11 | 1.44 | + | 1.89E-02 |
| Metabolism | 1968 | 301 | 238.87 | 1.26 | + | 4.52E-02 |

**Table 6:** Upregulated regulated TGA vs ctrl. Over-representation of GO biological processes in Upregulated TGA vs control

| GO biological process | Homo sapiens (REF) | Client Text Box Input |  |  |  |  |
| --- | --- | --- | --- | --- | --- | --- |
|  |  | # | expected | Fold Enrichment | +/- | P value |
| regulation of autophagy | 299 | 39 | 15.97 | 2.44 | + | 5.44E-03 |
| protein localization | 1927 | 150 | 102.93 | 1.46 | + | 2.29E-02 |
| phosphate-containing compound metabolic process | 2112 | 162 | 112.82 | 1.44 | + | 1.99E-02 |
| organelle organization | 3168 | 227 | 169.22 | 1.34 | + | 1.89E-02 |
| cellular protein modification process | 3158 | 225 | 168.69 | 1.33 | + | 3.15E-02 |

### Placental Nutrient Transporter expression

As our GoElite clustering analysis identified nutrient transporters as different between groups we investigated this further at the protein level. Staining of SLC38A2 (SNAT2), a transporter commonly associated with idiopathic intrauterine growth restriction, was substantially reduced in the syncytium of HLHS placentas compared to both control and TGA cases (Figure 1A-C). Members of the LAT family of transporters also demonstrated differential staining between control, HLHS and TGA placentas. SLC7A5 (LAT1) was present in the syncytium of all placentas but staining in the fetal endothelium was reduced in the HLHS placentas compared to both the control and TGA (Figure 1D-F). Similarly, in the HLHS placentas, SLC7A8 (LAT2) staining demonstrated reduced expression in both the syncytial microvillous membrane (MVM) and nuclei compared with control and TGA placentas (Figure 1G-I). The heavy chain of the LAT heterodimers SLC3A2 (CD98) was not differentially expressed in any of the placentas (data not shown).

**Figure 1:**
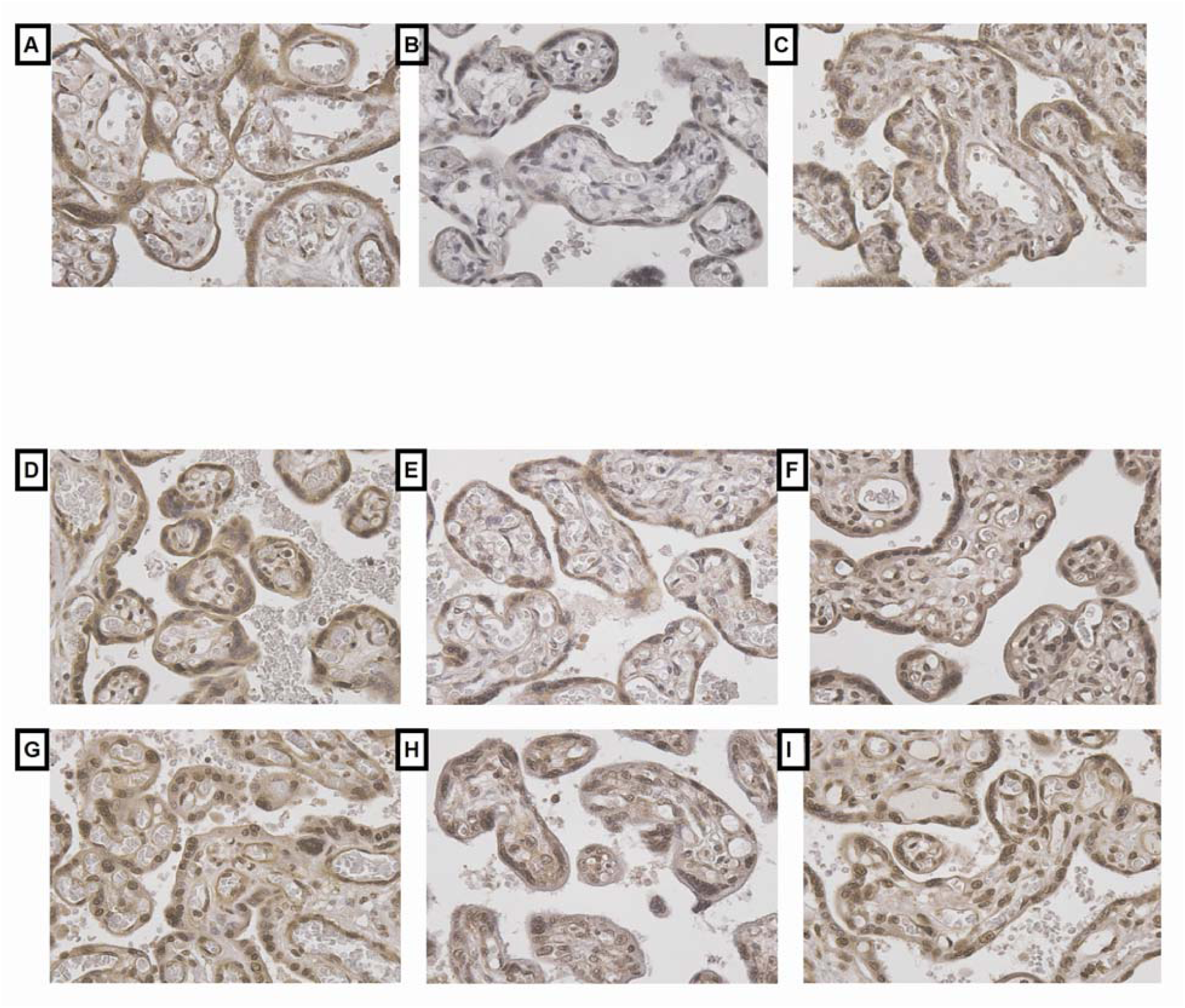
Detection of SNAT2 (A-C), LAT1 (D-F), and LAT2 (G-I) proteins by immunohistochemistry in control (A, D, G), HLHS (B, E, H) and TGA (C, F, I) term placentas. SNAT2 expression (associated with IUGR) is lower in HLHS placentas compared to control and TGA. LAT1 expression is reduced in fetal endothelial cells in HLHS versus control and TGA. HLHS placentas have lowered expression of LAT2 in syncytial microvillous membranes and nuclei compared to control and TGA. This demonstrates a lack of proper nutrient transport in HLHS fetuses, leading to lower placental and birth weights.

Expression of glucose transporter isoforms SLC2A1 (GLUT1) (Figure 2A-C) and SLC2A3 (GLUT3) (Figure 2D-F) demonstrated relocalization and altered expression. GLUT1 was present in both the MVM and basal membrane (BM) of the syncytium in the controls but appeared to be relocalized to the syncytial MVM in the HLHS placentas. Interestingly, stronger staining was seen in both MVM and BM in the TGA placentas compared to HLHS and control placentas. GLUT3 staining in the syncytium and fetal endothelium was reduced in the HLHS compared to both control and TGA placentas.

**Figure 2:**
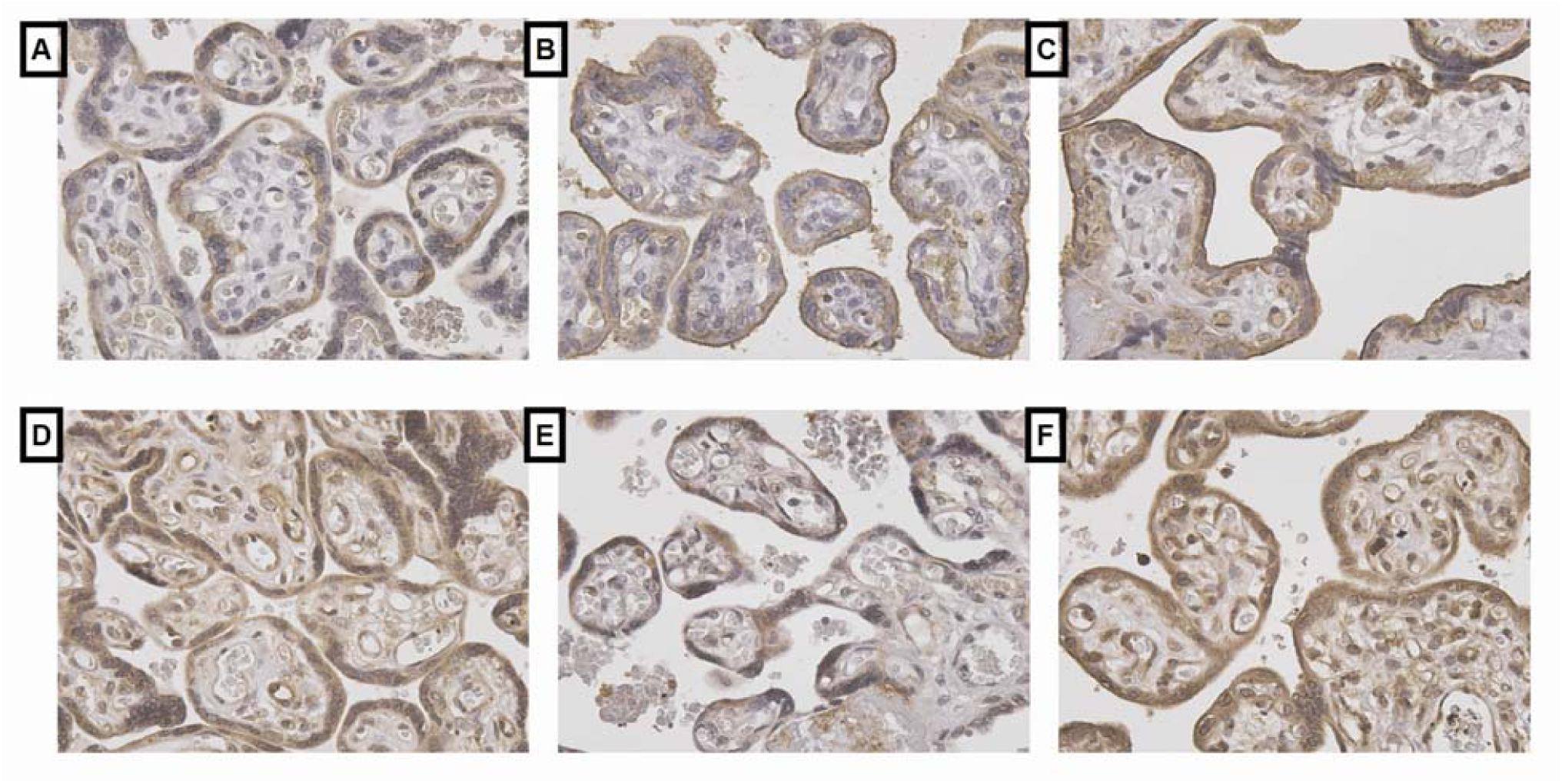
Detection of GLUT1 (A-C) and GLUT3 (D-F) proteins by immunohistochemistry in control (A, D), HLHS (B, E) and TGA (C, F) term placentas. GLUT1 expression has been relocalized to the syncytial Microvillous membrane (MVM) in HLHS placentas compared to control. Expression of GLUT1 appears higher in both the MVM and BM of TGA placentas compared to control. GLUT3 expression is reduced in syncytium and fetal endothelial cells in HLHS versus control and TGA. MVM: microvillous membrane; BM: basal membrane.

**Figure 3:**
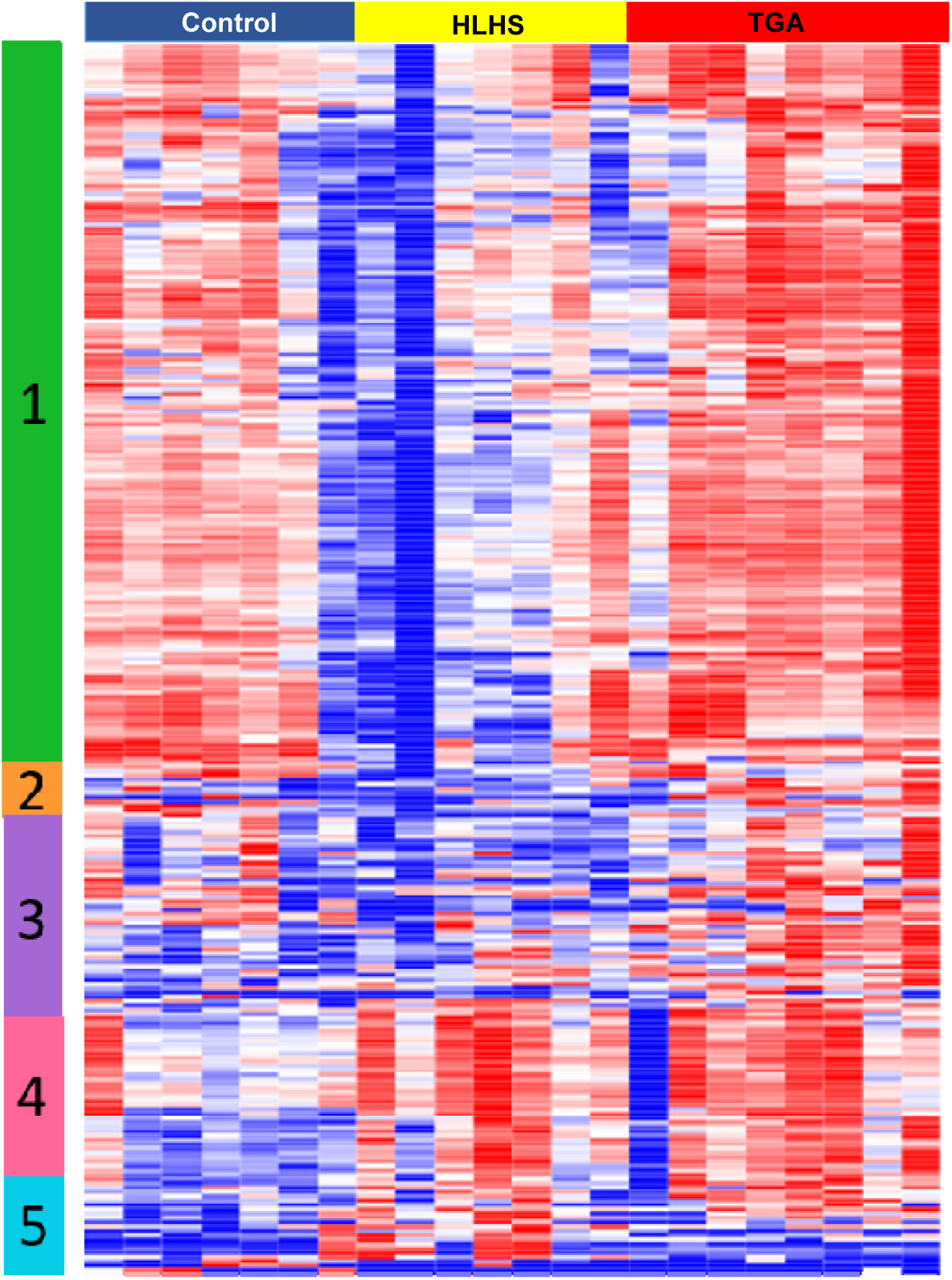
Unbiased gene clustering using GO ELITE from control (blue, n = 7), HLHS (yellow, n = 7), and TGA (red, n = 8) placentas. Cluster 1 (green) represents genes associated with left/right symmetry patterning, heart looping and muscle development, and energy synthesis. This cluster is significantly downregulated in the HLHS placentas compared to both control and TGA. Cluster 4 (pink) represents genes involved in pregnancy and hormone stimulation, which is generally upregulated in HLHS and TGA versus control. However, cluster 5 (teal) contains transporters, which are only upregulated in HLHS vs control and TGA. Individual gene lists are found in Supplemental Data.

## Discussion

The findings of the current study are the first to identify vascular and structural alterations in the villus tree of TGA placentas. We demonstrated intraparenchymal fibrin deposits and reductions in fetal vasculature at the maternal-fetal interface which may diminish the surface area available for nutrient and oxygen transport and impact transfer to the growing fetus. Together, these changes are consistent with placental dysmaturation and align with the phenotype of HLHS placentas as shown by Jones et al. [1]. The placental abnormalities are also associated with reduced head circumferences in both HLHS and TGA, however, consistent with other clinical observations from CHD birth cohorts (see references I placed in the intro), we found no differences in birth weight between TGA and controls. An increased ratio of birthweight to placental weight infers increased placental efficiency in the TGA cohort compared to controls and may represent one mechanism that underlies this difference in fetal growth. To investigate the underlying mechanisms associated with these more efficient placentas, we utilized RNA sequencing and immunohistochemistry.

Recently, epidemiological studies have identified differences in placental-related complications associated with different subtypes of CHD. Llurba et al.[22], reported that HLHS cases have a high incidence of SGA or IUGR but not preeclampsia, which was interpreted as meaning that placentation in these cases was normal. However, our previous [1] and current study identify significant placental abnormalities in HLHS cases which may represent the underlying mechanisms of the impaired fetal growth. We also identify common placental vascular disruption in two CHD subtypes that appear independent of the fetal growth phenotype.

Our pathway analysis of upregulated genes in HLHS vs control identified over-representation of genes involved in the Pantherdb biological process ‘female pregnancy.’ Upon further investigation, these genes include leptin, CRH, pregnancy-specific glycoproteins, and PAPP-A, factors which may represent a compensatory mechanism in response to the abnormal placental structure. We had previously demonstrated increased leptin RNA levels in HLHS Vs. term using qPCR [1]. Similar pathway findings and hypotheses have been postulated in twin-twin transfusion placentas [23]. Interestingly, these factors were not increased in the TGA placentas indicating they were not perturbed at the time of delivery. Moreover, despite the increase in transcription of these factors, the attempt at placental compensation by the HLHS placentas fails to achieve normal growth patterns. The failure of these compensatory mechanisms appears to be at several different levels. While our pathway analysis demonstrates an over-representation of ‘amino acid transport’ in the HLHS DE gene list, there is no increase in the expression of mTOR members RPTOR or RICTOR in response to the compensation factors suggesting no increase in transcription of nutrient transport regulatory mechanisms. However, there is down regulation of major cellular pathways such as Cell Cycle, Generation of second messenger molecules and Interferon Signaling. These significant perturbations to the placenta would impact its development, growth and function. Assessment of these pathways at the protein level, while beyond the current study, may illuminate further mechanisms. In contrast, pathway analysis of the DE genes upregulated in TGA compared to control or HLHS demonstrates a placenta with increased general transcription, metabolism, TCA cycle and protein localization capable of sustaining fetal growth despite the reduction in fetal vasculature. Interestingly, no pathways were over-represented in DE genes that were down-regulated in TGA placentas compared with control or HLHS.

Common anomalies in the placental vascular tree in the placentas from HLHS and TGA may represent disruptions in common developmental pathways in the placenta and heart in early gestation. Our transcriptome studies have highlighted several genes/pathways of interest that are implicated both in early trophoblast and cardiac development. For example, the IGF2 pathway; transcriptional levels of members of this pathway are reduced in this HLHS cohort compared with control (IGF2, IGFBP2, 3 & 5) and elevated in the TGA cohort compared with the HLHS (IGF2, IGF2BP1, IGFBP1, 2, 3 & 4), no differences are seen in between TGA and control groups. Disruptions in the IGF pathway are frequently associated with impaired placental development, growth and function and recently this pathway was identified as playing a significant role in cardiac development [24]. Disruption of this pathway would significantly impact both placental and heart development as well as placental function. Genes involved in left-right patterning/symmetry and heart looping were significantly lower in the HLHS placentas, again highlighting disruptions to common pathways involved in both organs development. Cited2 is required both for proper placental capillary patterning and left-right patterning of the body axis, and activates proliferation through the TGF-β pathway. Our transcriptome data shows Cited2 is downregulated in HLHS placentas, consistent with dysregulation of both the heart and placenta [25] [26].

Regulation of cellular processes and development occurs at multiple levels and it is possible that post-transcriptional changes might be triggered by altered protein activity rather than expression. For example, in spontaneous IUGR, reduced placental mTOR activity shifted SNAT2 trafficking towards proteasomal degradation, contributing to reduced fetal amino acid availability and restricted fetal growth [27]. Interestingly, while we do not see over-representation of pathways involved in proteasomal degradation in our transcriptome, our immunohistochemistry results do demonstrate a significant loss in amino acid transporter proteins, most impactful SNAT2, in the HLHS placentas. Furthermore, the protein expression of LAT1 (SLC7A5) does not reflect the increase seen in the transcriptional analysis. The lack of transporters leading to reduced nutrient transport in HLHS placentas would significantly contribute to the diminished fetal growth in HLHS patients, as many studies in humans and animal models of spontaneous IUGR have verified [28-30]. In contrast, the normal nutrient transporter expression in TGA placentas may contribute to why these fetuses achieve normal birth weights.

Despite the limitations of our current study, which include cohort size and the use of FFPE tissues which limit assessment of protein expression, we have identified multiple placental mechanisms that are disrupted in two subtypes of CHD both dependent and independent of fetal growth, highlighting the need for further in depth investigation into the placenta in congenital heart disease. Furthermore, this study is a “snapshot” look at the placenta at time of delivery and therefore the need for complementary investigations that study the pregnancy in longitudinal manner will be necessary.

This study identifies the disruption of multiple placental mechanisms in two subtypes of CHD. The HLHS placenta demonstrates reduced vascular development with an insufficient compensatory upregulation of transporter activity. The TGA placenta demonstrates similar vascular development abnormalities but greater placental efficiency as noted by the ratio of placental: fetal weight and transporter expression. Transcriptome findings provide further insights on the molecular mechanisms that link fetal cardiac and placental development. Such insights may ultimately lead to targeted therapies that improve fetal growth and, in turn, postnatal CHD outcomes.

